# Extracellular symbiont colonizes insect during embryo development

**DOI:** 10.1101/2023.12.04.569952

**Authors:** Miguel Ángel González Porras, Inès Pons, Marleny García-Lozano, Shounak Jagdale, Christiane Emmerich, Benjamin Weiss, Hassan Salem

## Abstract

Insects typically acquire their beneficial microbes early in development. Endosymbionts housed intracellularly are commonly integrated during oogenesis or embryogenesis, whereas extracellular microbes are only known to be acquired after hatching by immature instars such as larvae or nymphs. Here, however, we report on an extracellular symbiont that colonises its host during embryo development. Tortoise leaf beetles (Chrysomelidae: Cassidinae) host their digestive bacterial symbiont *Stammera* extracellularly within foregut symbiotic organs, and in ovary-associated glands to ensure its vertical transmission. We outline the initial stages of symbiont colonization and observe that while the foregut symbiotic organs develop three days prior to larval emergence, they remain empty until the final 24 hours of embryo development. Infection by *Stammera* occurs during that timeframe, and prior to hatching. By experimentally manipulating symbiont availability to embryos in the egg, we describe a 12-hour developmental window governing colonization by *Stammera*. Symbiotic organs form normally in aposymbiotic larvae, demonstrating that these *Stammera*-bearing structures develop autonomously. In adults, the foregut symbiotic organs are already colonized following metamorphosis and host a stable *Stammera* population to facilitate folivory. The ovary-associated glands, however, initially lack *Stammera*. Symbiont abundance subsequently increases within these transmission organs, thereby ensuring sufficient titers at the onset of oviposition ∼29 days following metamorphosis. Collectively, our findings reveal that *Stammera* colonization precedes larval emergence, where its proliferation is eventually decoupled in adult beetles to match the nutritional and reproductive requirements of its host.

## Introduction

Insects evolved a remarkable diversity of specialized cells and organs to house and faithfully propagate beneficial microbes [1–4]. Correspondingly, symbionts vary in how they colonize and populate these structures [5], often reflecting their beneficial role relative to host development and metabolic requirements.

A number of insect taxa harbor their symbionts intracellularly within specialized cells known as bacteriocytes [6–9]. Bacteriocytes are often colonized by a monoclonal population of microbes [6,10], but can also host multiple symbiont strains that are metabolically distinct [11]. Most intracellular symbionts are vertically transmitted during embryogenesis or oogenesis [9,12–15], reflecting a high degree of integration between symbiont and host [16]. For example, aphids transmit their nutritional endosymbiont *Buchnera* during embryo development through calibrated cycles of exocytosis and endocytosis [17]. *Buchnera* cells released from maternal symbiotic organs colonize cells fated to become bacteriocytes in the developing embryo [17]. Symbionts can even leverage the host’s developmental machinery to facilitate colonization, as demonstrated in carpenter ants and the intertwined regulatory network shared with their endosymbiont *Blochmannia* [18].

For extracellular symbionts residing along the gut lumen [19,20], within specialized crypts [21–24], or on cuticular surfaces [25,26], symbiont colonization is only demonstrated to take place following embryo development [27], i.e. after hatching [28], and during immature developmental stages such as larvae or nymphs. For example, bean bugs (*Riptortus pedestris*) acquire their beneficial *Caballeronia* symbionts from the environment every generation [29]. *Caballeronia* colonizes its host during a specific developmental window after hatching [22], triggering the rapid formation of specialized symbiont-harboring gut crypts [30]. Maternal secretions can also ensure the strict vertical transmission of extracellular symbionts in newly-hatched insects, as demonstrated in wasps [31], beetles [32,33], and numerous stinkbugs [21,34,35]. In this study, however, we report that infection by an extracellular symbiont can precede eclosion from the egg, by describing the colonization dynamics of a beneficial microbe in tortoise beetles (Chrysomelidae: Cassidinae).

Tortoise beetles are hosts to *Candidatus* Stammera capleta, a γ-proteobacterial symbiont [36–40]. *Stammera* is housed extracellularly in specialized organs near the foregut where it upgrades the digestive physiology of its host by supplementing pectinases and other plant cell wall-degrading enzymes [36,37,39,41]. In adult females, *Stammera* is also maintained in ovary-associated glands to ensure the microbe’s vertical transmission [36]. Cassidines propagate the symbiosis by depositing a symbiont-bearing ‘caplet’ at the anterior pole of each egg (Figure 1, a, b) [36,41]. The caplets are populated by ∼12 spherical secretions where *Stammera* is embedded (Figure 1, c-e) [36,41]. Developing embryos are separated from *Stammera* by a thin membrane (Figure 1, e) that remains intact until the final 24 hours of embryogenesis [41]. Experimental removal of the caplet disrupts *Stammera* transmission, yielding symbiont-free (aposymbiotic) larvae that exhibit a diminished digestive capacity and low survivorship [36,41]. Reintroducing symbiont-bearing caplets to aposymbiotic cassidines after hatching does not rescue infection, suggesting that *Stammera* colonizes its host during embryo development [41], in contrast to the post-hatch acquisition routes described for extracellular insect symbionts [27].

**Figure 1:**
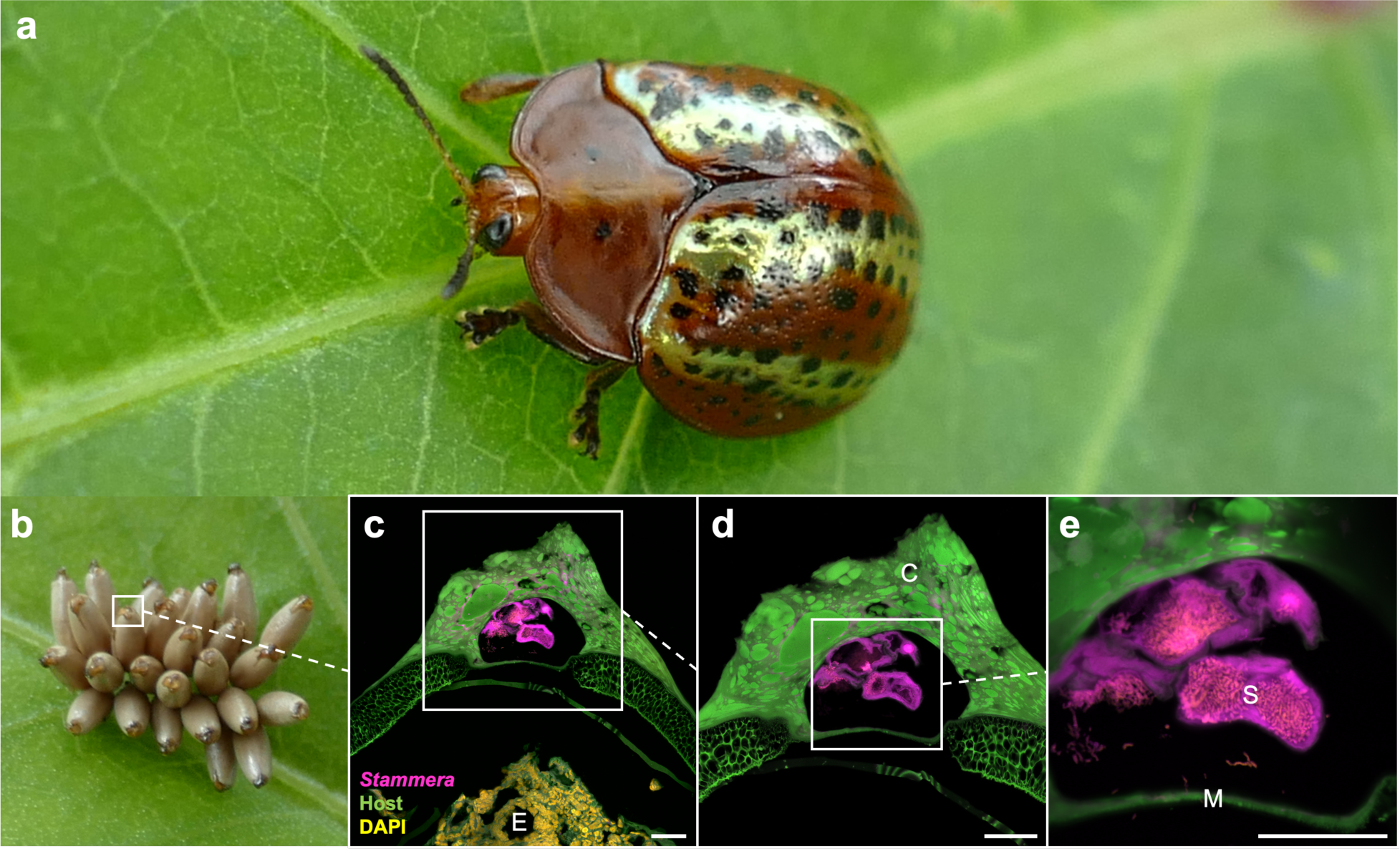
Tortoise beetles transmit *Stammera* via egg caplets. **(a)** The tortoise beetle *Chelymorpha alternans.* **(b)** Eggs deposited on an *Ipomoea batatas* leaf, each topped with a *Stammera*-bearing caplet at the anterior pole. **(c-e)** Fluorescence *in situ* hybridization (FISH) on longitudinal section of the egg caplet, where *Stammera* is separated from the developing embryo via a thin membrane. Probes: *Stammera* 16S (magenta), beetle 18S (green), and DAPI-stained DNA (yellow). Abbreviations: C, caplet; M, caplet membrane; E, embryo; S, *Stammera*-bearing spheres. Scalebar is included for reference: 50 µm.

Here, we (**i**) determine the colonization dynamics of *Stammera* relative to the early developmental stages of the tortoise beetle *Chelymorpha alternans*, (**ii**) define a narrow temporal window for symbiont acquisition following experimental manipulation, (**iii**) test whether the formation of symbiotic organs depends on symbiont presence, and (**iv**) quantify the proliferation of *Stammera* after metamorphosis relative to the nutritional and reproductive requirements of its beetle host.

## Results and discussion

### *Stammera* colonizes its host during embryo development

To investigate the morphogenesis of the foregut symbiotic organs and their colonization by *Stammera*, we applied fluorescence *in situ* hybridization (FISH) using cross sections of embryos dissected from eggs (96, 72, 48, 24 hours prior to hatching), along with larvae spanning the five instar stages of *C. alternans* (Figure 2). We observe that evaginations resulting in the foregut symbiotic organs begin to form in the final 72 hours of embryogenesis, and that these structures become fully developed at the −48-hour mark (Figure 2). However, the foregut symbiotic organs remain empty until the last day of embryo development, during which *Stammera* colonization takes place (Figure 2). Our current findings confirm that infection by *Stammera* occurs prior to larval eclosion, in contrast to the post-hatch dynamics described for other extracellular insect symbionts [27]. Across stinkbugs [21,23,29,34,35], wasps [31], ants [25], bees [20], and beetles [26], hatchlings are initially aposymbiotic but eventually acquire their extracellular microbes from the environment, through trophallaxis, or by consuming maternal secretions [27]

**Figure 2:**
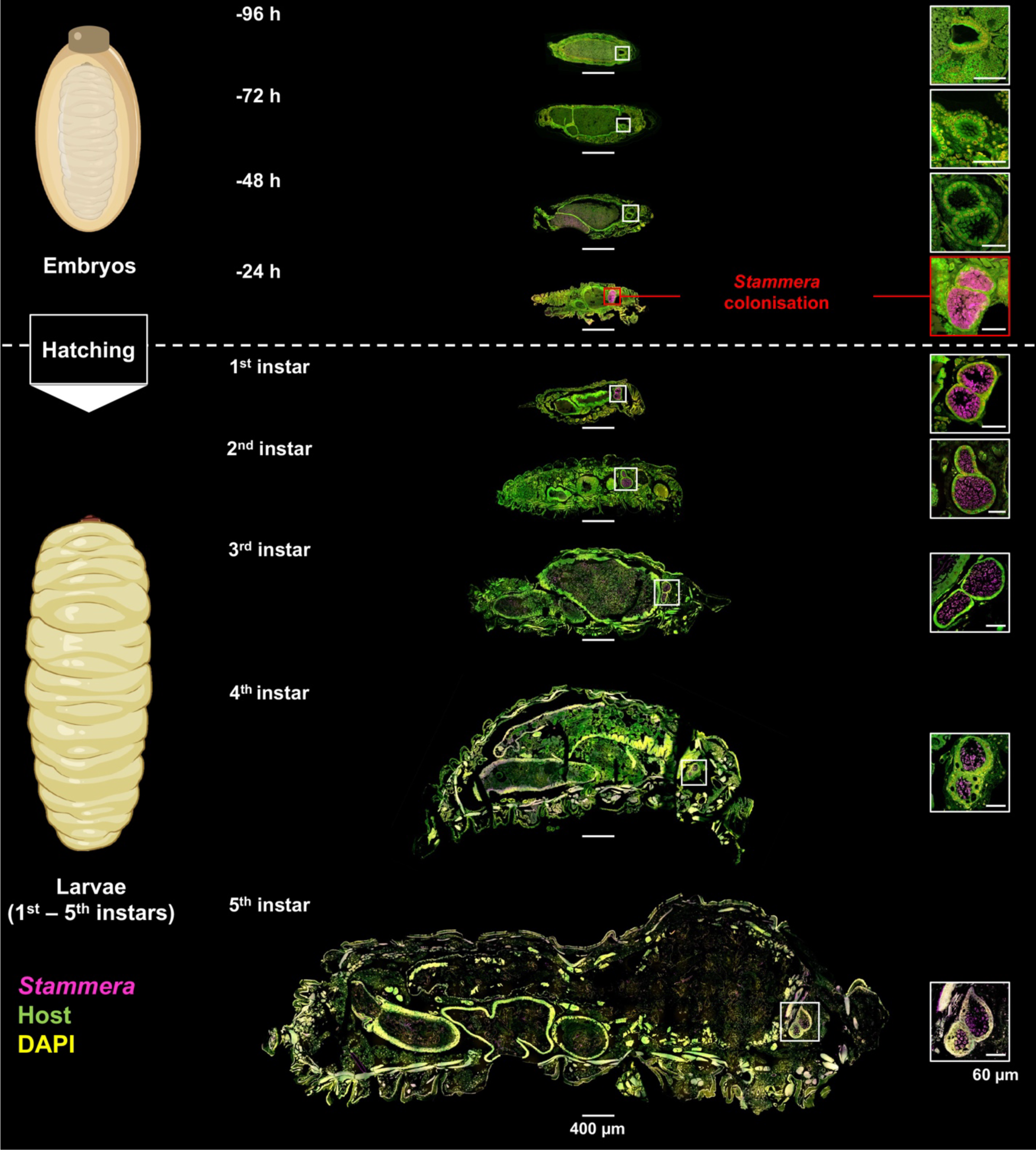
*Stammera* colonization dynamics during embryo and larval development. FISH on sagittal sections of dissected *C. alternans* embryos (96, 72, 48, and 24 hours prior to hatching) and larvae (1^st^ – 5^th^ instars). Insets show the foregut symbiotic organs at a greater magnification. The foregut symbiotic organs begin to form between 72 and 48 hours prior to larval eclosion, and are colonized by *Stammera* within 24 hours. Probes: *Stammera* 16S rRNA (magenta), host 18S rRNA (green), and DAPI-stained DNA (yellow). Scalebars are included for reference: 60 and 400 µm.

*Stammera*’s colonization dynamics may reflect its host’s requirements for pectinases and other plant cell wall-degrading enzymes upon larval eclosion [36,41]. We observe that newly emerged larvae shift away from their chorions and onto the leaf surface within 43.1 (± 4.82) minutes after hatching. While speculative, the immediate onset of folivory in larvae may select for the foregut symbiotic organs to be fully developed and colonized prior to hatching (Figure 2). Stage-specific host requirements could explain the divergent colonization dynamics described for *Stammera* relative to other extracellular symbionts, such as *Tachikawaea* in urostylidid bugs [21] and *Burkholderia* in darkling beetles [26]. Urostylidid nymphs initially consume maternally-provisioned, nutritionally-rich jelly ahead of eventually transitioning to sap-feeding weeks later [21]. As *Tachikawaea* supplements essential nutrients to balance sap-based diet, its post-hatch proliferation within specialized gut crypts in nymphs coincides with the commencement of plant-feeding, thereby matching the nutritional requirements of the host [21]. Similarly, in the darkling beetle *Lagria villosa*, infection by *Burkholderia* ectosymbionts takes place after hatching [26], reflecting the microbe’s role in upgrading the defensive biochemistry of its host during molting and, eventually, metamorphosis [42,43]. The dorsal symbiotic organs of developing embryos are initially symbiont-free. However, these structures are later colonized in hatching larvae [26], contrasting early histological descriptions by Hans-Jürgen Stammer that suggested a pre-hatch infection in the congeneric *Lagria hirta* [44].

### A pre-hatch window for symbiont colonization

The mechanisms guiding symbiont colonization are often tightly regulated and highly synchronized relative to insect development [5,45]. Governing that interplay are cellular [9], morphological [26,46], and behavioral adaptations [35] to ensure symbiont uptake while mitigating the risk of secondary exposure to pathogens and parasites [19]. Infection competence can thus vary throughout the lifespan of an insect, resulting in a defined window for symbiont colonization [22,25,26]. Bean bugs, for example, acquire their *Caballeronia* symbionts from the environment every generation [29], but do so more faithfully during their 2^nd^ and 3^rd^ nymphal instar stages relative to the 1^st^, 4^th^ and 5^th^ [22]. For darkling beetles, larvae are efficiently colonized by their defensive ectosymbionts after hatching, but are less likely to acquire their microbes when exposed at a later stage [26]. Here, we explored whether similar developmental constraints govern *Stammera* colonization in tortoise beetles.

Guiding our experimental framework were two observations: (i) foregut symbiotic organs are developed and colonized by *Stammera* 24 hours prior to larval eclosion (Figure 2), and (ii) aposymbiotic insects do not reacquire the microbe after hatching [41]. This suggests that access to *Stammera* at least 24 hours prior to hatching is critical for successful colonization and that infection efficiency decreases over time. Therefore, we reapplied *Stammera*-bearing spheres to the anterior pole of caplet-free eggs at two timepoints: 24 and 12 hours prior to hatching. In addition to a control group where egg caplets were left untreated, we compared *Stammera* infection frequencies across all treatments in 2^nd^ instar larvae (Figure 3).

**Figure 3:**
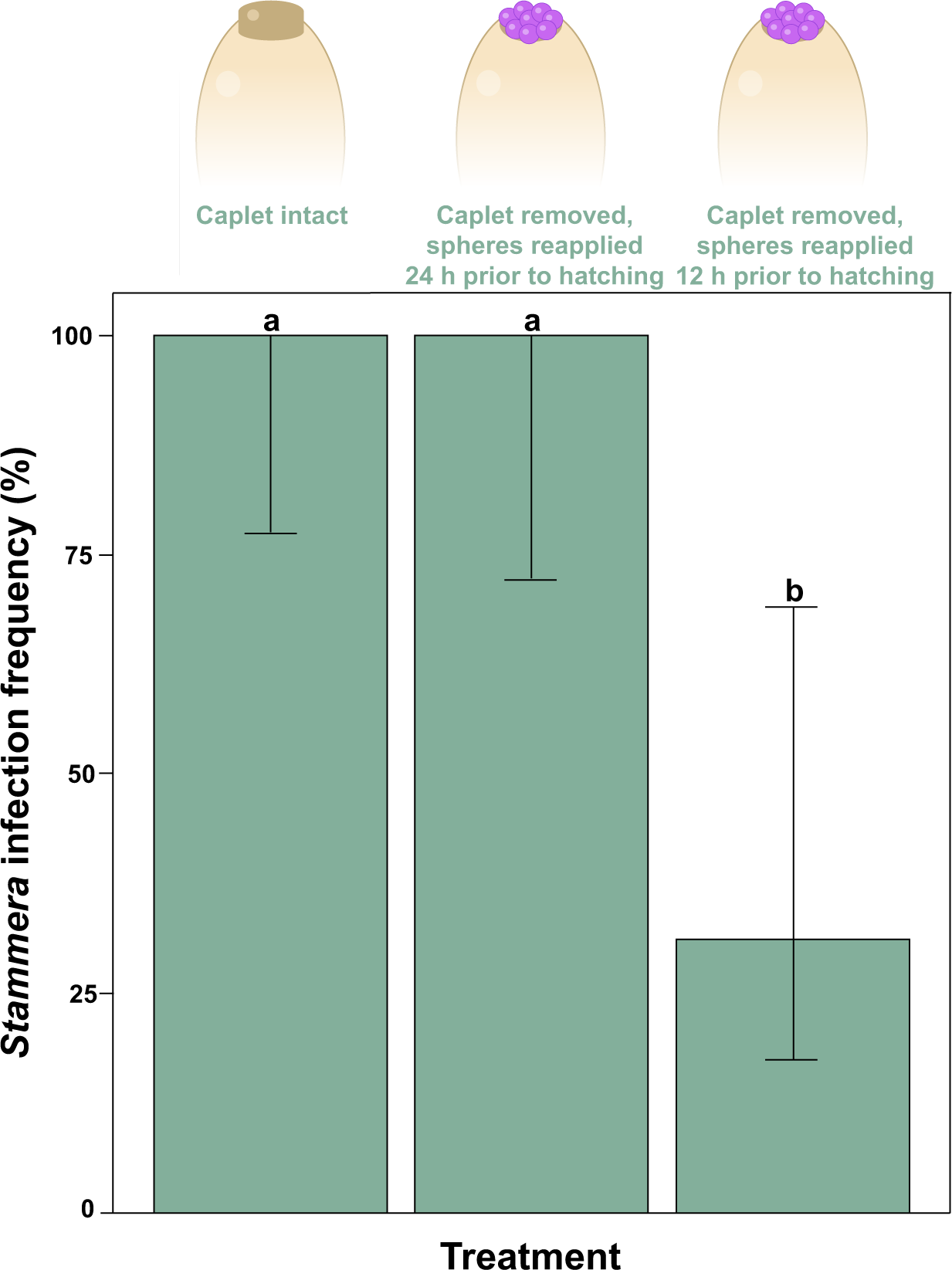
A narrow developmental window for symbiont colonization. *Stammera* infection frequencies in *C. alternans* larvae following experimental manipulation of the egg caplet and its symbiont-bearing spheres (Fisher’s exact test, *p* < 0.001). Number of samples = 33 larvae; Caplet intact (13), *Stammera*-bearing spheres reapplied 24h prior to hatching (10), *Stammera*-bearing spheres reapplied 12 h prior to hatching (10). Whiskers denote the 95% binomial confidence intervals. Different letters above the bars indicate significant differences.

Reestablishing embryo access to *Stammera*-bearing spheres 24 hours prior to eclosion restores symbiont infection rates to levels mirroring the untreated control group (Fisher’s exact test, *p* = 1) (Figure 3). This is consistent with the timing of caplet membrane rupture [41] and the onset of symbiont colonization in the foregut symbiotic organs (Figure 2). In contrast, *Stammera* was acquired less efficiently when spheres were resupplied 12 hours later (Fisher’s exact test, *p* = 0.011). Delayed access constrained symbiont infection competence (Figure 3), pointing to a narrow developmental window for efficient colonization. Several factors may underpin this process [5,46,47], including mechanical adaptations ensuring that the symbiotic organs are only populated by *Stammera* as opposed to environmental microbes, or potential pathogens encountered after hatching. In bean bugs, for example, midgut crypts colonized by *Caballeronia* become irreversibly sealed following morphological modifications to a connective ‘sorting’ channel [46,47]. We observe a similar duct connecting the foregut to the symbiotic organs in tortoise beetles [36]. While speculative, this channel may become less permeable during the latter stages of embryo development, thereby limiting microbial passage over time.

### Foregut symbiotic organs develop independently of *Stammera* infection

Symbiotic organs vary considerably in their morphology and developmental features [4]. Many reflect ancient evolutionary origins where organ formation and development proceeds autonomously; whereas others are triggered by microbial factors that promote cellular differentiation and morphogenesis in the host [48].

Here, we clarified whether *Stammera* induces the development of the foregut symbiotic organs in its beetle host. Two observations indicated that these derived ceca develop independently, despite an obligate co-dependence [36,41] and an intertwined co-evolutionary history between *Stammera* and tortoise beetles [37,39]. First, the foregut symbiotic organs form prior to symbiont colonization (Figure 2), highlighting that the differentiation process precedes contact with *Stammera*. Second, the evolutionary loss of *Stammera* in a subset of Cassidinae species does not correspond to the absence of symbiotic organs [39]. These beetles retain vestigial structures devoid of microbes, implying that *Stammera* does not trigger their formation [39]. To test this experimentally, we compared the morphology of the foregut symbiotic organs in symbiotic and aposymbiotic larvae. These organs developed fully in both groups (Figure 4), retaining their sac-like morphology in aposymbiotic and symbiotic larvae alike (Figure 4, a-d). Additionally, the epithelial extensions where *Stammera* is typically embedded were also present, highlighting the conserved cellular features of the symbiont-bearing structures in an aposymbiotic state (Figure 4, e-h).

**Figure 4:**
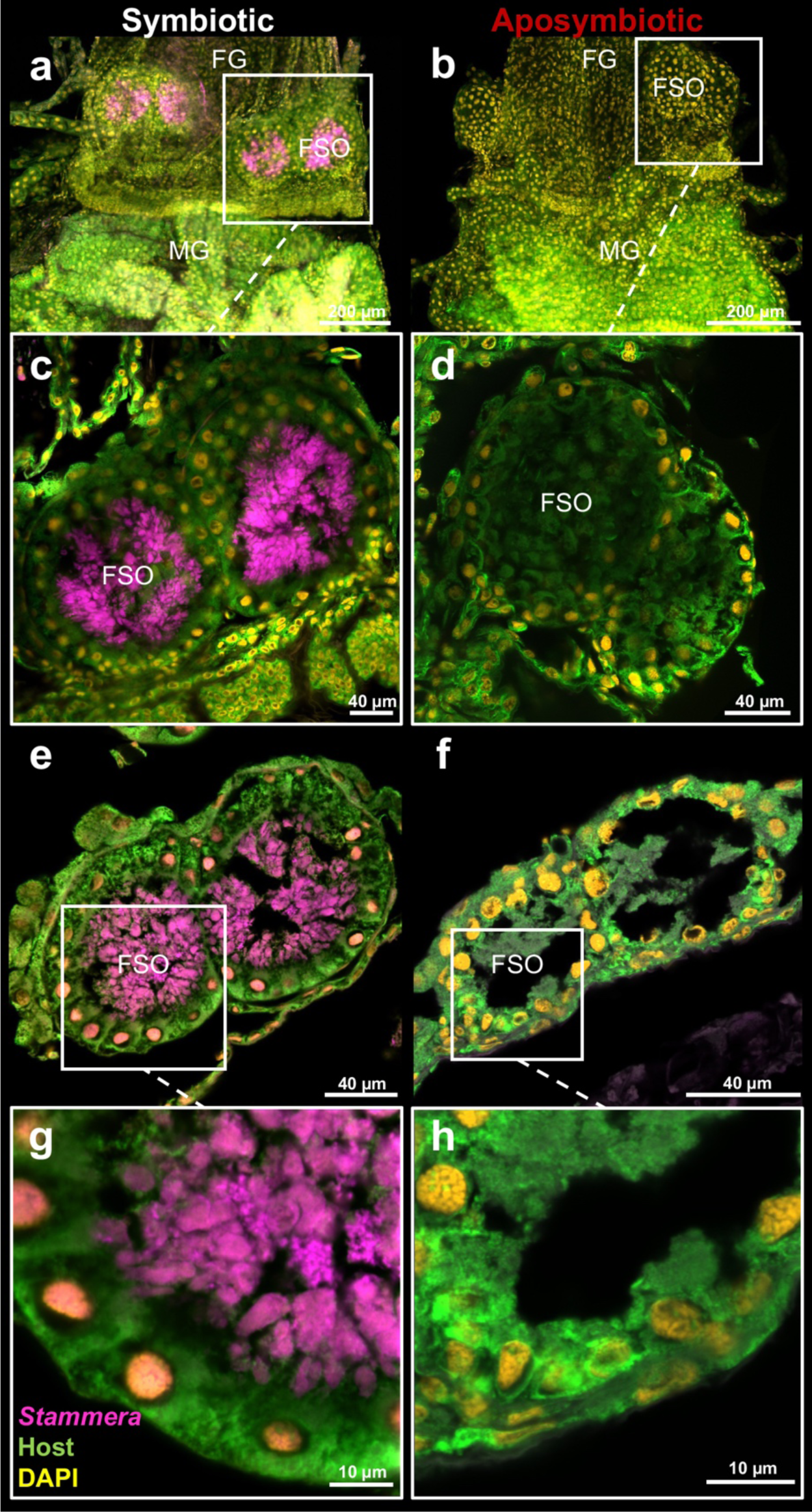
The foregut symbiotic organs develop independently of *Stammera*. FISH micrographs of whole-mount (**a-d**) and cross-sections (**e-h**) of the foregut-midgut tract dissected from symbiotic and aposymbiotic 2^nd^ instar larvae. Insets show the foregut symbiotic organs at a greater magnification. Probes: *Stammera* 16S rRNA (magenta), host 18S rRNA (green), and DAPI-stained DNA (yellow). Abbreviations: FG, foregut; MG, midgut; FSO, foregut symbiotic organ. Scalebars are included for reference.

While the developmental origin of the foregut symbiotic organs remains undescribed in tortoise beetles, it is possible that it represents an ingrained process that was coopted to house and maintain beneficial microbes in the gut. Other leaf beetles engaging in functionally convergent digestive symbioses also host their microbial partners in sac-like structures derived from gastric ceca [49–52]. This is analogous to stinkbugs and their diverse nutritional symbioses with extracellular bacteria [27]. Stinkbugs belonging to the Plataspidae [23,53], Urostylididae [21], Acanthosomatidae [54], Alydidae [22,29,55], and Pentatomidae [56–59] families all house their symbionts in crypts developing in the posterior midgut. Recent findings indicate that the underlying molecular and cellular processes appear to be decoupled from symbiont presence, at least in a subset of species [57]. In the case of the pentatomid *Plautia stali*, the experimental loss of its obligate *Pantoea* symbiont does not alter crypt formation and cellular differentiation [57]; an observation that is consistent with our finding that the foregut symbiotic organs also develop independently of *Stammera* in tortoise beetles (Figure 4). This contrasts diverse symbiotic systems where microbial colonization induces organ formation [4], as demonstrated in *Caballeronia*-harboring bean bugs [30], *Vibrio*-hosting squids [60–62], and leguminous plants in partnership with rhizobia [63,64].

### *Stammera* population dynamics relative to adult nutritional and reproductive requirements

Symbiont density can drastically fluctuate throughout the life cycle of its host [4,65]. Such differences can be especially pronounced in partnerships where the microbial partner is housed in specialized cells or organs [65–67]. Beyond maximizing the benefits derived through symbiosis, these organs enable the host to regulate symbiont abundance relative to its own development and metabolic requirements [4,45]. That dynamic is evident in weevils in their symbiosis with *Sodalis*, a nutritional symbiont that contributes to cuticle synthesis in its host by supplementing tyrosine and phenylalanine [6]. In the first days after metamorphosis, symbiont abundance sharply increases within midgut-associated bacteriomes, matching its host’s requirements for aromatic amino acids that are required for exoskeleton development [65]. Upon cuticle formation, however, *Sodalis* is rapidly eliminated through host-driven apoptosis and autophagy [65]. Most strikingly, the symbiont continues to persist within apical bacteriomes associated with the ovaries, thereby ensuring its vertical transmission [65]. Here, we explored the proliferation of *Stammera* upon adult eclosion, and in light of the its discrete localization within two functionally divergent symbiotic organs: foregut symbiotic organs to facilitate folivory (Figure 5, a), and ovary-associated glands to ensure the symbiont’s vertical propagation (Figure 5, b) [36].

**Figure 5:**
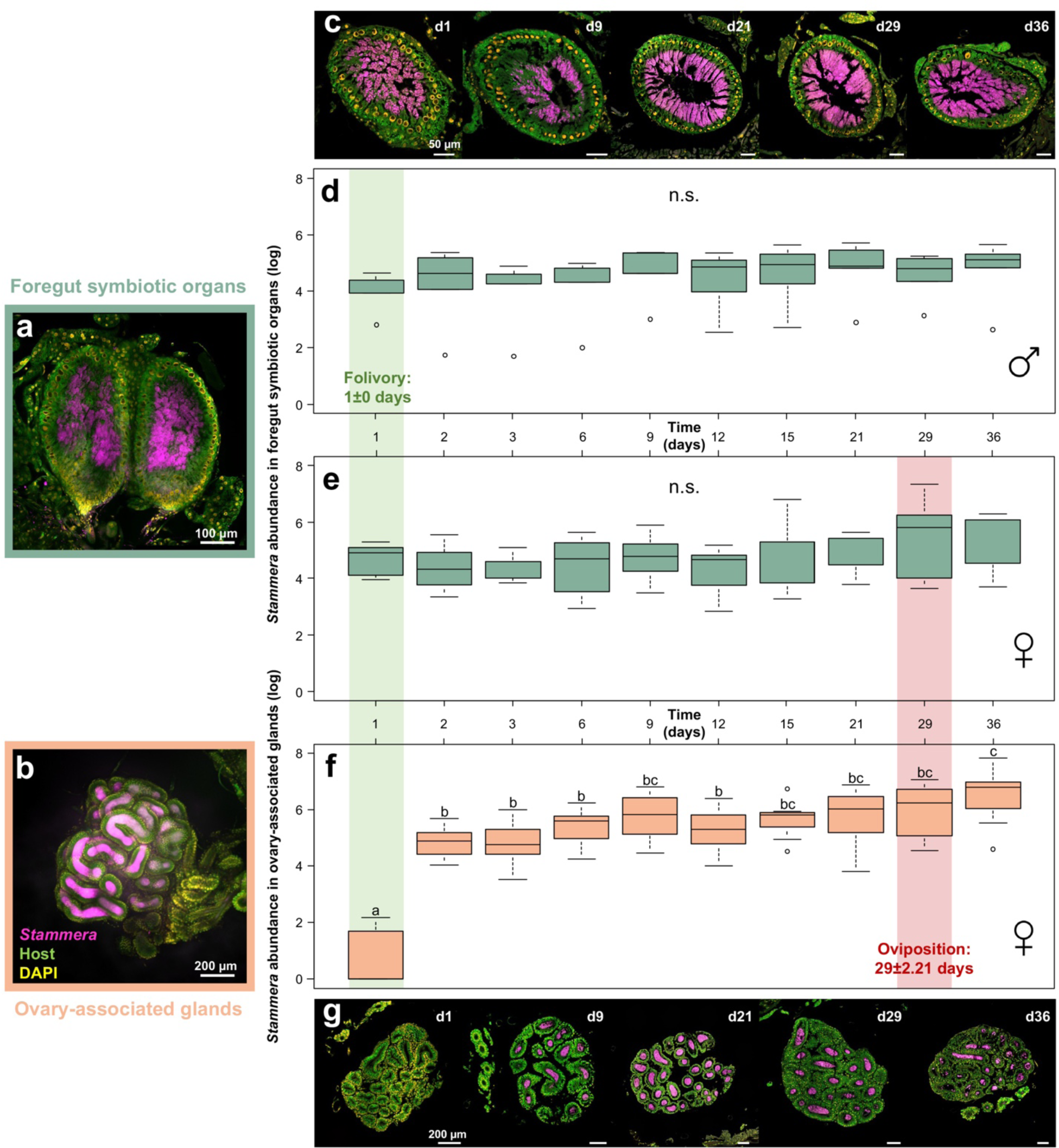
Symbiont proliferation relative to folivory and oviposition in adult beetles. (**a**) FISH micrographs on whole-mounts of foregut symbiotic organs, and (**b**) ovary-associated glands. **c**, FISH cross-sections of foregut symbiotic organs 1, 9, 21, 29 and 36 days following metamorphosis. *Stammera* abundance in the foregut symbiotic organs in (**d**) males (LM, F_9,40_= 0.8, *p* = 0.62), (**e**) females (LM, F_9,60_= 1.01, *p* = 0.38), and in (**f**) the ovary-associated glands of (LM, F_9,54=_ 19.91, *p* < 0.001) based on the quantification of the symbiont’s 16S rRNA gene copy numbers. Lines represent medians, boxes indicate 25–75 percentiles, and whiskers denote range. Different letters above boxes indicate significant differences. The green- and red-faded frames denote the onset of folivory and oviposition, respectively. **(g)**, FISH micrographs of ovary-associated glands 1, 9, 21, 29 and 36 days following metamorphosis. Probes: *Stammera* 16S rRNA (magenta), host 18S rRNA (green), and DAPI-stained DNA (yellow). Scalebars are included for reference. Abbreviations: n.s., not significant; d, day.

We observe that the foregut symbiotic organs are already populated by *Stammera* following metamorphosis in both males and females (Figure 5 c-e). This corresponds with the immediate resumption of folivory (1±0 days), highlighting the host’s metabolic requirements for symbiosis-derived digestive enzymes [36,41]. How *Stammera* subsists within the foregut symbiotic organs, despite the likely immune challenges and epithelial transformation that accompanies its host’s metamorphosis [68], is still unknown. It is possible that these organs undergo a similar morphological and spatial reorganization as observed in *Sodalis*-harboring weevils [6], thereby ensuring the persistence of symbiosis during pupation. In quantifying symbiont abundance within the foregut symbiotic organs (Figure 5, d, e), we observe a highly stable *Stammera* population during adulthood (Males: LM, F_9,40_= 0.8, *p* = 0.62; Females: LM, F_9,60_= 1.01, *p* = 0.38), matching the steady feeding behavior recorded for cassidines [69], including *C. alternans* [70,71]. For each sampling day, males and females did not differ in their *Stammera* titers (LMM, F_9,90_= 0.7, *p* = 0.71), which is consistent with the conserved morphology of the foregut symbiotic organs across sexes [36,39].

For the ovary-associated glands (Figure 5, b), *Stammera* does not colonize these organs immediately after metamorphosis (Figure 5, f, g). Of the seven females examined on the first day of adulthood, four lacked the symbiont in their transmission glands (Figure 5, g). We observe that symbiont proliferation differed significantly within the ovary-associated glands (LM, F_9,54=_ 19.91, *p* < 0.001) (Figure 5, f, g), in contrast to the stable *Stammera* population within the foregut symbiotic organs (Figure 5, c-e; Males: LM, F_9,40_= 0.8, *p* = 0.62; Females: LM, F_9,60_= 1.01, *p* = 0.38). A significant peak is observed on the second day, followed by a more gradual increase before oviposition (Figure 5, f, g). The diverging symbiont proliferation dynamics in both organs appears to reflect their distinct functions relative to the maintenance and propagation of *Stammera*. Folivory resumes following metamorphosis and continues throughout adulthood [70,71], requiring the foregut symbiotic organs to be already colonized by *Stammera* (Figure 5, c), and for the symbiont population to persist at a relatively stable level (Figure 5, d, e). In contrast, we observed that egg-laying commences 29±2.21 days following adult eclosion. The temporal lag between folivory and the onset of oviposition matches our observation that the ovary associated glands become densely occupied later in adulthood (Figure 5, f, g). This revealed that *Stammera*’s population in adult females is decoupled across two types of symbiotic organs and appears to be regulated to meet the nutritional requirements and reproductive cycle of its host (Figure 5).

Our quantification of *Stammera* titers in adults focused on young, reproductively active beetles (Figure 5). How senescence and diapause impact symbiont density is also of interest and worthy of exploring in future studies [65–67,72]. Tortoise beetles experimentally induced to enter diapause cease feeding and pause egg-laying [70]. As similar observations are noted for senescing cassidines [69], it is possible that these insects modulate their symbiont titers, as shown in aphids [66], weevils [65], and ants [67]. For example, older aphids recycle *Buchnera* through Rab7 recruitment and lysosomal activity following bacteriocyte cell death [66]. Clarifying the population dynamics of *Stammera* during the latter developmental stages of its host can shed light on the mechanisms by which extracellular symbionts are regulated, and, potentially, recycled upon fulfilling their host-beneficial roles.

### Conclusions and outlook

By describing the colonization dynamics of *Stammera capleta* within its beetle host, we uncovered a pre-hatch route that is uncommon for extracellular insect symbionts. Several open questions remain, including (i) how *Stammera* contends with the likely transformation of its habitat during metamorphosis by its host, (ii) which molecular and cellular mechanisms underlie the morphogenesis of symbiotic organs during embryo development in cassidines, and (iii) whether these factors reflect a shared evolutionary origin with other symbiotic leaf beetles, including members of the Eumolpinae [50] and Donaciinae subfamilies [49]. Given recent advances in microdissections, transcriptome sequencing and RNA interference, our future efforts will complement a growing set of studies on the developmental basis and regulation of symbiotic organs and bacteriomes [18,30,57], and extend our knowledge on the adaptations ensuring the maintenance of specialized microbes in the gut [73].

## Methodology

### Insect rearing

A laboratory culture of *Chelymorpha alternans* is continuously maintained at the Max Planck Institute for Biology in Tübingen, Germany. The insects are reared in mesh cages (60 x 60 x 90 cm) along with their host plant, *Ipomoea batatas* [74]. Eggs were reared in an incubator (Memmert, Germany) at a constant temperature of 26°C to control for embryo development as previously reported by Pons *et al* [41].

### Fluorescence *in situ* hybridization

To localize *Stammera* in the symbiotic organs of *C. alternans* at different developmental stages, we applied fluorescence *in situ* hybridization (FISH) on tissue sections and whole-mounts. We designed an oligonucleotide probe specifically targeting the 16S rRNA sequence of *Stammera* from *C. alternans*, SAL227 (5’GGTCTTGAAAAAAAAAGATCCCC’3) using the software ARB [75]. We included the eukaryotic 18S rRNA probe EUK-1195 (5’GGGCATCACAGACCTG’3) [76] to localize *C. alternans* cells. All probes were dually labelled with fluorescent dyes at their 5’ and 3’ ends. Unless specified, fixation was done in 4% formaldehyde in PBS for 4 h at room temperature under gentle shaking (400 rpm). We visualized the samples using a LSM 780 confocal microscope (Zeiss, Germany) and an Axio Imager Z1 Microscope (Zeiss, Germany).

### Preparation of Technovit sections

Embryos and larvae were embedded and sectioned in Technovit. Due to the fragility of early embryos, whole eggs were placed into Carnoy’s solution (Ethanol: chloroform: acetic acid = 6: 3: 1) and incubated overnight at room temperature for fixation [77] before washing and dissection from chorion in 70% Ethanol. Subsequently they were dehydrated in a series of increasing concentrations of ethanol: 3x 80%, 3x 90%, 3x 96%, and 3x 100% (10 min each), followed by three incubations in 100% acetone for 15 min each. From the embryo stage 24 h before hatching onwards, these steps were adjusted due to a different sample size and composition. These were fixed in 4% paraformaldehyde (paraformaldehyde: PBS 1X) (v/v) at room temperature, shaking at 500 rpm for 4-10 h, depending on their size. For more efficient penetration by the fixative, larvae appendages, peripheral chaeta, and furcal chaeta were removed one hour after the start of fixation. Fixation of 4^th^ and 5^th^ instar larvae was interrupted after the first half of the incubation time. These larvae were cut in half and subjected to two incubations in chloroform at room temperature for 24 h under shaking (800 rpm) before returning them to the fixation solution. After fixation, larvae were dehydrated in a series of increasing concentrations of tertiary butanol in water (v/v) as follows: 3x 80% (30 min each), 1x 90% (1 h), 1x 96% (1 h), 3x 100% (2 h each), followed by two incubations in 100% acetone for one hour each. Dehydration steps were performed under shaking (800 rpm) at 26°C (100% tertiary butanol). Following dehydration, all samples were embedded in Technovit 8100 following the manufacturer’s protocol and clustered in Teflon molds (Kulzer, Germany). The Technovit-embedded samples were sagittal-sectioned at 7 µm using either home-made glass knives (embryos and 1^st^ – 3^rd^ instar larvae) or metal HistoBlades (4^th^ and 5^th^ instar larvae) (Kulzer, Germany) on a conventional microtome (Leica, Germany). Sections were transferred to water droplets on HistoBond glass slides (Marienfeld, Germany) kept at 50°C over a warm plate for 20 min to promote section unfolding. FISH was performed as described [78].

### FISH on paraffin sections

Foregut symbiotic organs of symbiotic and aposymbiotic 2^nd^ instar larvae and foregut symbiotic organs and ovary-associated glands of adult females at different timepoints following metamorphosis were dissected and fixed. Dehydration was achieved by an increasing ethanol series of 60, 70, 80, 96 and 100% (v/v) for one step of 1 h for 60, 70 and 80% ethanol and three steps of 1 h each for 96 and 100% ethanol. After dehydration, samples were gradually transferred into paraffin by passing through three incubations of Roti-Hostol (CarlRoth, Germany) at room temperature (2x 40 min, 1 x overnight), followed by incubations at 60°C in Roti-Histol:paraffin (1:1 v/v) for 60 minutes and paraffin (Paraplast High Melt, Leica, Germany) (3 x 60 min, 1 x overnight). The paraffin-embedded samples were cross sectioned at 10 µm using a conventional microtome and mounted on poly-L-lysine-coated glass slides (Epredia, Germany) using a 40 °C water bath. Paraffin sections were dried at room temperature overnight and incubated at 60°C for 1 h to improve tissue adherence. The sections were dewaxed with Roti^®^-Histol in three consecutive steps for 10 min each followed by ethanol 100% for 10 min. Next, slides were dried at 37°C for 30 min. Probes were dissolved at 900 nM in the hybridization buffer containing 35% formamide (v/v), 900 mM NaCl, 20 mM Tris-HCl pH 7.8, 1% blocking reagent for nucleic acids (v/v) (Roche, Switzerland), 0.02 SDS (v/v) and 10% dextran sulfate (w/v). Hybridization was done at 46°C for 4 h. Sections were rinsed in 48°C washing buffer (70 mM NaCl, 20 mM Tris-HCl pH 7.8, 5 mM EDTA pH 8.0, and 0.01% SDS (v/v)) and transferred to fresh 48°C washing buffer for 15 min followed by room temperature washes in PBS (20 min) and milliQ water (1 min). Sections were counterstained with DAPI (5 µg/ml) for 10 min at room temperature, dipped in milliQ water, dipped in ethanol 100% and dried at 37°C for 30 min. Slides were mounted using the ProLong^®^ Gold antifade mounting media (Thermo Fisher Scientific, MA, USA), cured overnight at room temperature, and stored at −20°C until visualization.

### Whole-mount FISH

Whole-mount FISH was performed on symbiotic and aposymbiotic 2^nd^ instar larvae as well as adult females. Fixed samples were washed in PBS at room temperature under gentle shaking for 30 min and permeabilized in acetic acid 70% (v/v) at 60°C for 1 min. Samples were washed three times in PBS at room for 5 min. Samples were carefully laid on KIMTECHScience precision wipes to remove PBS from the samples. For hybridization, samples were transferred in hybridization buffer (0.9 M NaCl, 0.02 M Tris/HCl pH 8.0, 0.01% SDS) containing 900 nM each probe and 5 µg/ml DAPI for DNA counterstaining. Samples were hybridized at 46°C for 4 h and transferred to 48°C washing buffer (0.07 M NaCl, 0.02 M Tris/HCl pH 8.0, 0.01% SDS, 5 mM EDTA) for 15’. Samples were mounted on microscopy slides with VectaShield mounting media (Vector, Burlingame, CA, USA).

### Experimental elimination of *Stammera capleta*

Three egg masses (∼30 eggs each) were collected from three different *C. alternans* females. Each was then separated into two experimental treatments, an untreated control and an aposymbiotic treatment. To generate aposymbiotic larvae, caplets were removed from eggs using sterile dissection scissors, followed by surface sterilization with 99% ethanol as previously outlined [41]. Experimental treatments were maintained as described above. 2^nd^ instar larvae were collected ten days after hatching. Foregut-symbiotic organs (FG-SO) of larvae were dissected and fixed in 4% formaldehyde/PBS (v/v) (Electron Microscopy Sciences, PA, USA) at room temperature 4 h under shaking (500 rpm). Samples were stored in PBS:Ethanol (0.5X:50%) at −20°C until microscopy processing.

### Experimental manipulation to elucidate the timing of symbiont acquisition

Three egg masses originating from different *C. alternans* females were collected. Each mass was then separated into three experimental treatments: (a) untreated control and (b) eggs whose caplets were removed, and *Stammera*-containing spheres were resupplied 24h prior to hatching, and (c) eggs whose caplets were removed and *Stammera*-containing spheres were manually resupplied 12h prior to hatching. Across both time points, fresh spheres were carefully extracted and resupplied to caplet-free eggs as previously described by Pons et al [41]. Three days after hatching, DNA was extracted from larvae using the EZNA^®^ Insect DNA Kit and *Stammera* infection frequencies of each treatment were evaluated by *Stammera*-specific diagnostic PCR, as previously described in [41]. Diagnostic PCR was conducted on an Analytik Jena Biometra TAdvanced Thermal Cycler (Analytik Jena AG, Germany) using a final volume of 20 μl containing 1 μl of DNA template, 0.5 μM of each primer and 2x DreamTaq Green PCR Master Mix (ThermoFisher Scientific, MA, USA). The following cycle parameters were used: 5 min at 95°C, followed by 34 cycles of 95°C for 30 s, 57.7 or 62°C (depending on the primer) for 30 s, 72°C for 1 min and a final extension time of 2 min at 72°C [41]. Primers used for diagnostic PCR are listed in Table S1.

### Folivory and oviposition monitoring in adult beetles

To determine when larvae transition away from their chorions and onto their host plants after hatching, we monitored the commencement of folivory in *C. alternans* across five egg masses and recorded the first instance of leaf damage by eclosing larvae. To time the resumption of folivory and record the onset of oviposition in *C. alternans* adults, we monitored beetles in small mesh cages (30 x 30 x 30 cm) supplemented with a single host plant. A total of 6 mating pairs were placed in 6 separate cages immediately following metamorphosis. Feeding and oviposition were monitored daily by direct observation of foliar damage and presence of egg clutches, respectively.

### *Stammera* population dynamics in adult beetles

*Stammera* population dynamics within the foregut symbiotic organs and the ovary-associated glands were determined using qPCR. Seven and five egg clutches were collected from females and males, respectively, and sibling groups were maintained on individual small mesh containers (30 x 30 x 30 cm) with a host plant until they reached adulthood. A single female and male were sampled per replicate and their foregut symbiotic organs (females and males) and ovary-associated glands (females) were dissected at days 1, 2, 3, 6, 9, 12, 15, 21, 29 and 36 following metamorphosis, accounting for 140 female and 50 male samples, respectively. Following dissection, symbiotic organs were preserved in 500 µL of 100% ethanol and kept at −70°C until DNA extraction. DNA was extracted from *C. alternans* symbiotic organs using the EZNA^®^ Insect DNA Kit, and *Stammera* relative abundance was estimated using an Analytik Jena qTOWER^3^ cycler (Analytik Jena AG, Germany). The final reaction volume of 25 μl included the following components: 1 μl of DNA template, 2.5 μl of each primer (10 μM) (Table S1), 6.5 μl of autoclaved distilled H_2_O, and 12.5 μl of Qiagen SYBR Green Mix (Qiagen, Germany). Primer specificity was verified *in silico* by comparison with reference bacterial sequences in the Ribosomal Database and NCBI. Additionally, PCR products were sequenced to confirm primer specificity *in vitro*. Standard curves (10-fold dilution series from 10^−2^ to 10^−8^ ng·μl^−1^) were generated using purified PCR products and measuring their DNA concentration using a NanoDrop TM1000 spectrophotometer. The following cycle parameters were used: 95°C for 10 min, followed by 45 cycles of 95°C for 30 s, 62.7°C for 20 s, and a melting curve analysis was conducted by increasing temperature from 60 to 95°C during 30 s. Based on the standard curve, absolute copy numbers were calculated, which were then used to extrapolate symbiont relative abundance by accounting for the single copy of the 16S gene in *Stammera*’s genome, as previously described [41].

### Statistical analyses

*Stammera* population dynamics within the foregut symbiotic organ and ovary-associated glands throughout female development was analyzed using general linear models (LMs) after reverse transformation and validation of a normal distribution (Table S2a). The time and replicate variables were used as fixed factors. The *Stammera* population dynamics within the foregut symbiotic organ along male development was also evaluated using a general linear model (LM) after reverse transformation and validation of a normal distribution and using time and replicate as fixed factors (Table S2a). After statistical modeling, Tukey’s HSD pairwise comparisons were performed using the ‘glht’ function with Bonferroni corrections. To determine whether there was an effect of sex on the *Stammera* population dynamics within the foregut symbiotic organ of *C*.

*alternans*, a general linear mixed model (LMM) was performed after reverse transformation and validation of a normal distribution, using time, sex, and their interactions as fixed factors (Table S2b). In addition, the replicate variable was considered as a random factor because females and males were not harvested from the same egg clutches. Fisher’s exact test was used to assess the effect of experimental manipulation on *Stammera* infection frequency in *C. alternans* larvae (Table S2c). Statistical analyses were performed using the software R version 3.5.3 (R Core Team, 2019) [79], using the multcomp package for Tukey’s HSD pairwise comparisons [80], nlme package for LMM [81], and ggplot2 package for boxplot visualization [82].

## Acknowledgements

We thank Christine Henzler and Alejandra Leyva for technical assistance, Martin Kaltenpoth, Aileen Berasategui, and members of the Electron Microscopy and BioOptics Facilities for helpful comments during project development. Funding from the Max Planck Society, Humboldt Foundation (to IP), and the German Research Foundation (to HS; SA 3105/2-1) is gratefully acknowledged.

## Author contributions

MAGP, IP and HS conceived the project and designed the experiments. IP, MAGP, SJ and CE performed histological processing pipelines. MAGP, IP, CE, and SJ carried out microscopy experiments. IP and MGL produced aposymbiotic insects. MAGP and IP performed bioassays. IP, MAGP and MGL performed qPCR experiments. IP conducted the statistical analyses. MAGP, IP and HS wrote the manuscript with input from all coauthors. IP and HS secured funding. HS supervised the project.

